# An embryonic system to assess Wnt transcriptional targets

**DOI:** 10.1101/129056

**Authors:** Jahnavi Suresh, Nathan Harmston, Ka Keat Lim, Prameet Kaur, Helen Jingshu Jin, Jay B. Lusk, Enrico Petretto, Nicholas S. Tolwinski

## Abstract

During animal development, complex signals determine and organize a vast number of tissues using a very small number of signal transduction pathways. These developmental signaling pathways determine cell fates through a coordinated transcriptional response that remains poorly understood. The Wnt pathway is involved in a variety of these cellular functions, and its signals are transmitted in part through a β-catenin/TCF transcriptional complex. Here we report an *in vivo Drosophila* assay that can be used to distinguish between activation, de-repression and repression of transcriptional responses, separating upstream and downstream pathway activation and canonical/non-canonical Wnt signals in embryos. We find specific sets of genes downstream of both β-catenin and TCF with an additional group of genes regulated by Wnt, while the non-canonical Wnt4 regulates a separate cohort of genes. We correlate transcriptional changes with phenotypic outcomes of cell differentiation and embryo size, showing our model can be used to characterize developmental signaling compartmentalization *in vivo*.

## Introduction

Signaling pathways elicit cellular responses in part by regulating the transcription of specific cohorts of target genes. Signaling pathways that are crucial for development, homeostasis and tumorigenesis have both negative (repressive) and positive (activation and de-repression) effects on transcription ^1^. Negatively regulated targets can have different biological activities from positively regulated targets ^2^. The Wnt signaling pathway provides a striking example of this phenomenon where Wnt signals can elicit a variety of cellular responses including differentiation, growth, and polarity ^3-6^. Increased Wnt signaling has apparently opposite roles on cell proliferation: excessive Wnt signaling leads to over proliferation of cancer cells, but it is also required to maintain undifferentiated, quiescent stem cells ^7,8^. As a result, therapeutic interventions that block Wnt in tumours are likely to have both beneficial and detrimental consequences; when cancer growth is inhibited, useful stem cells are likely to be lost as a side-effect ^9,10^. It is probable that these opposing effects occur through the transcriptional activation of different Wnt target genes, raising the intriguing possibility that therapies targeted downstream could avoid the detrimental effects of disrupting the whole pathway.

Wnt signalling and its dysregulation has been implicated in a variety of developmental disorders and diseases, including diabetes, Robinow Syndrome, cancer and aging ^11-13^. Wnt regulation appears to be highly context-specific, affecting different genes in different cell types at different developmental stages ^14^, and the features defining how a Wnt target gene is regulated are not fully understood ^15^. Wnt signaling refers to a series of signaling pathways or networks divided into non-canonical and canonical. Non-canonical signaling is a collection of signal transduction pathways that do not use TCF/β-catenin for their transcriptional outputs ^16^. These are associated with planar or apical-basal polarity affecting pathways or calcium signaling. In vertebrates, these are driven by non-canonical Wnts-4, -5a, -5b, -6, 7a, -7b, and -11. In *Drosophila*, the main non-canonical or Planar Cell Polarity (PCP) pathway uses Wnt indirectly ^17^, but there are two characterized Wnts that do not signal through the canonical pathway. Wnt5 binds to the receptor Ryk and mediates axonal pathfinding ^18^, while Wnt4 functions through PTK7 in opposition to canonical Wnt1 ^19,20^ to regulate polarity, cell migration and invasion ^21^.

The canonical Wnt signaling pathway is largely mediated via β-catenin binding to the transcription factor TCF ^4,5^. TCF binds to DNA through a consensus sequence, CCTTTGATCTT, at genes it activates in conjunction with β-catenin, and at the same site with Groucho/TLE at genes it represses. Genes at Gro/TCF sites can be de-repressed by Wnt signaling, where Gro and β-catenin competitively bind to TCF setting up a switch where β-catenin can remove Gro/TCF leading to derepression, or replace Gro/TCF with β-catenin/TCF leading to activation ^22^. Additionally, genes can be actively repressed upon signaling activation through TCF/β-catenin binding to a novel consensus site, AGAWAW ^23^, or through a second repressor Coop ^24^. Specificity can be increased through helper sites bound by the C-clamp region ^15,23,25-30^. In vertebrates, the four TCF gene family members function as both transcriptional repressors and activators ^31^. *Drosophila* has only one gene that encodes TCF, which must perform both functions, making this system simpler to manipulate genetically. This feature that was recently used to study activation and derepression of Wnt targets in *Drosophila* tissue culture cells lacking TCF ^32^. In studies of TCF in fly embryos, the difference between the two functions of TCF becomes apparent when loss of function mutants are compared to dominant negative TCF transgenes ^33^. Loss of function embryos show a loss of patterning, but the embryos remain large ^33^. In contrast, expression of dominant negative TCF leads to a small embryo that lacks patterning ^5^. This finding led us to propose that the two roles of TCF might be separable at the transcriptional level, and led us to develop tools to analyse transcription *in vivo*.

We focus on developing methods for assessing transcriptional programs downstream of Wnt signalling, and identifying the processes and mechanisms involved. To this aim, we developed a naïve embryo system in which we can activate or repress different forms of Wnt signaling at various levels in the different pathways. This system allowed us to dissect the effects of Wnt signalling at both the phenotypic level of the whole organism and at the molecular level.

## Results

### Development of a naïve-embryo transcriptional assay system

We developed a transcriptional assay using the *Drosophila* embryo, where by using simple genetic manipulations we can create relatively naïve, homogeneous populations of cells, therefore minimizing the confounding effect of non-specific, secondary, and multiple signaling pathway effects that are often observed in gene expression studies ^34^. In normal *Drosophila* development, eggs are provided with maternal patterning signals. These signals include anterior-posterior patterning molecules such as Bicoid and Nanos, terminal patterning determinants such as Torso and Torsolike (EGF pathway related), and dorsal-ventral signals such as Toll and Dpp (NFκB and TGFβ signaling pathways) ^35-37^. These patterning signals determine the axes of the developing embryo and activate further signals that lead to specific cellular identities for each cell in the embryo. Removal of these basic patterning signals leads to eggs that develop a simple, un-patterned epithelium “naïve embryos”. For anterior-posterior patterning, we used a triple mutant (*bicoid, nanos, torsolike*) that eliminates anterior, posterior and terminal patterning respectively leading to highly compromised development (Supplementary Movies 1 and 2) ^38,39^. A further advantage of this system lies in the fact that these are maternal effect mutations, allowing the use of homozygous females that lay 100% mutant eggs, therefore removing the difficulty of identifying mutant embryos and avoiding the use of complicated germline clone techniques ^33,40,41^. This experimental setup creates a condition where all the embryonic cells are identical with respect to Wnt signalling.

### Basic phenotypic analysis of Wnt signaling

We introduced several genetic changes targeting specific components of the Wnt signalling pathway and assessed their consequences via a simple phenotypic assay (Fig. 1). This was accomplished by examining both the size of the embryos (normal/small) and their differentiation status (naked or with denticles) (Fig. 1C). Wg overexpression (Wg++) was accomplished by using the GAL4/UAS system to establish the hyper-activated pathway condition ^42^. The opposite condition was the expression of a dominant-negative allele (tcfΔN-short). This form of TCF lacks the Arm binding region, the Groucho binding sequence (GBS) and the C-clamp that are required for β-catenin binding, Gro repressor binding, and additional target specificity respectively (Supplemental Figure 1) ^15,26^. This makes tcfΔN-short a strong dominant negative form of TCF as it is unable to release DNA, interact with Gro or recruit the transactivating domain of β-catenin effectively blocking both activation and de-repression and leading to small, un-patterned embryos (Fig. 1) ^3^. We were unable to generate the perfect intermediate condition, or *tcf* maternally and zygotically mutant embryos as the genetics were too complicated, instead we overexpressed a longer form of tcfΔN (tcfΔN-long, S1). This TCF gene lacks only the N-terminal β-catenin binding domain and retains the GBS and C-clamp maintaining Gro dependent repression and de-repression, but lacks β-catenin dependent activation. This form of TCF does not respond to β-catenin and phenocopies loss of TCF in embryos (Fig. 1, and S1) ^33^.

**Figure 1:**
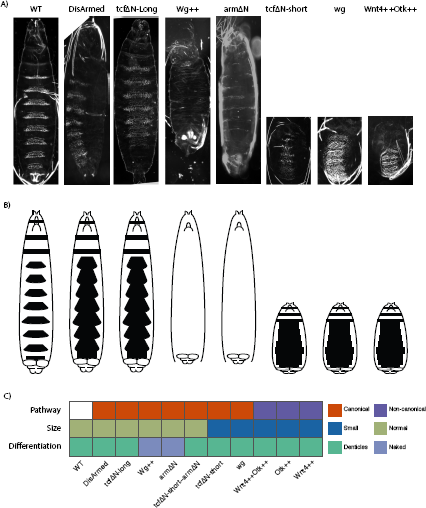
Mutations affecting components of the Wnt signalling pathway lead to various developmental phenotypes. (A) Phenotypes of the Wnt signaling conditions analysed, with large denticle covered embryos (DisArmed, tcfΔN-long), large naked embryos (Wg++, armΔN), and small denticle covered embryos (tcfΔN-short, *wg*, Wnt4++/Otk++) compared to wildtype (WT). (B) Schematic view of embryo size and denticle coverage as shown in black. (C) Embryos were classified based on their observed phenotype in terms of size (normal/small) and differentiation status (assessed by denticle coverage; covered/naked).

The classic *wingless* phenotype in *Drosophila* embryos shows a small denticle covered embryo ^43^. Similar phenotypes were observed for other strong loss of function alleles of Wnt signaling genes such as *arm*/β-catenin and *dishevelled* ^44,45^. tcfΔN-long expressing embryos, which lack transcriptional activation in response to Wnt activation, are large ^33^, whereas tcfΔN-short embryos, which lack activation in response to Wnts but retain repression of Wnt targets, are small ^5^. Under both conditions, patterning and the cell-fate decisions are disrupted in the same way (all epidermal cells make denticles), which suggests that transcriptional activation is required for differentiation and cell-fate determination, while regulation of transcriptional repression is required for cell proliferation and embryo size (Fig. 1).

Another way to establish the intermediate phenotype (large, un-patterned embryos) was the expression of a dominant negative version of Arm (DisArmed) where the transactivating region of the C-terminus is deleted along with a mutation in the Pygopus binding site ^23^. Expression of DisArmed blocks signaling by binding to TCF but not forming any activating complexes as transcriptional machinery is not recruited. We observed that these embryos showed a patterning phenotype where all cells made denticles, but the embryos were still large similar to loss of TCF (Fig. 1). This suggests that DisArmed blocks transcriptional activation, but still allows de-repression similar to TCF mutants ^33^.

In order to understand the effects of perturbing the non-canonical Wnt pathway, we used the non-canonical Wnt4 ligand that functions in opposition to the canonical Wg ^19,20^. We have recently shown that uniform expression functions in conjunction with the co-receptor PTK7 (Protein Tyrosine Kinase 7, *Drosophila* Offtrack or Otk) to oppose canonical signals in *Xenopus* and *Drosophila,* resulting in small un-patterned embryos similar to tcfΔN-short and *wg* ^19^. As Wnt4 opposes Wg, we hypothesized that this would provide a non-canonical, not functioning through *arm*/β-catenin and TCF, readout.

### Expression profiling reveals distinct transcriptional programs reflecting observed phenotypes

We examined changes in gene expression in all of the conditions using microarrays (see Methods). Even in our developmentally restricted system we identified 1,360 genes whose expression was significantly altered upon perturbing either canonical or non-canonical Wnt signalling (False Discovery Rate (FDR) < 1%). Hierarchical clustering of these genes revealed distinct expression patterns associated with perturbations affecting these separate branches of the Wnt signalling pathway (Fig. 2A, S2A). Expression of DisArmed showed a mild phenotypic change compared to WT, and this was mirrored in the similarity of their expression profiles (Fig. 1A). Overall, the transcriptional changes segregated according to whether the genetic perturbation was affecting the canonical (armΔN, tcfΔN-long, tcfΔN-short or Wg++), or non-canonical signalling (Wnt4++, Otk++). Full results of differential expression analyses and associated enrichments for biological processes and pathways are reported in Suppl. Table 1 and 2 respectively.

**Figure 2:**
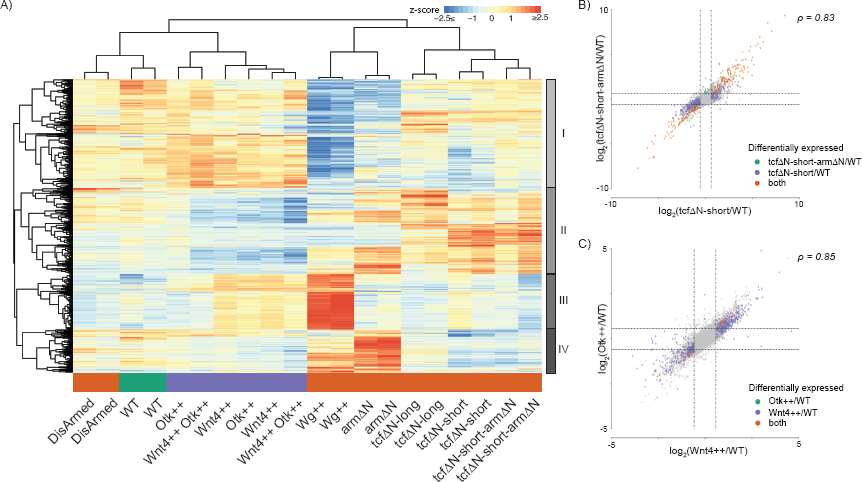
Transcriptional profiling identifies patterns of gene expression reflecting differences in canonical and non-canonical Wnt signalling. (A) Hierarchical clustering of gene expression profiles (z-scores) revealed a well-defined clustering of samples by branch of the Wnt signalling pathway perturbed, and the identification of four distinct clusters of genes with differing patterns of response. (B) Comparison of fold changes observed comparing tcfΔN-short against WT versus tcfΔN-short-armΔN against WT. Both are highly correlated indicating that they are located within the same branch of the Wnt signalling pathway (i.e. canonical signalling). Genes are highlighted depending on whether they are differentially expressed (absolute fold change > 1.5, FDR < 10%) in both conditions or only in one. (C) Comparison of fold changes observed comparing Wnt4++ against WT versus Otk++ against WT. Both are highly correlating indicating that they are located within the same branch of the Wnt signalling pathway (i.e. non-canonical signalling). Genes are highlighted depending on whether they are differentially expressed (absolute fold change > 1.5, FDR < 10%) in both conditions or only in one.

Within the canonical pathway, Arm primarily signals through TCF ^19^, as such the expression profiles of tcfΔN-short-armΔN and tcfΔN-short are highly similar (Fig. 2A, S2A) and show highly correlated changes in gene expression compared to WT (Fig. 2B), illustrating that these are epistatic. Wnt4 has been reported as primarily signalling via PTK7/Otk ^19^. In keeping with this, we found that overexpression of Wnt4 and Otk had highly correlated transcriptional responses compared to WT (Fig. 2A, 2C), providing further evidence of their functional interaction, epistasis and location within the same branch of the Wnt signalling pathway.

Across all conditions we identified four major clusters of genes (I-IV, Fig. 2A, S2B-C), each of which was associated with distinct enrichments for biological processes, pathways and transcription factor binding sites (TFBSs) (Fig. 3). Cluster I showed reduced expression upon stimulation of canonical Wnt signalling via Wg++ or armΔN and increased expression upon overexpression of the non-canonical branch (Fig. S2D). This cluster was enriched for processes associated with development, differentiation, cell-cell communication and morphogenesis (Figure 3A), and contained several transcription factors (TFs). The enrichment for Trl and TCF motifs suggests that cluster I may contain direct targets of canonical Wnt-mediated repression (Figure 3E). Genes present in cluster II are downregulated by overexpression of Wnt4 or Otk (Fig. 2A, S2D) and hence represent a set of genes that are putatively repressed by non-canonical Wnt signalling, but that also show upregulation upon the loss of the ability of Wnt to derepress genes repressed by TCF. Intriguingly, this cluster was enriched for glutathione metabolism genes (Fig. 3B), as well as for binding sites of the known Wnt target *Dll* ^46^ (Fig. 3E). Recent studies have discovered genes that are activated downstream of canonical Wnts, but do so independently of β-catenin through the Wnt/STOP pathway ^47^. Cluster III corresponded to a set of 256 genes that were upregulated by Wg++ but not by armΔN. This cluster was enriched for genes involved in processes and pathways relating to cell cycle and proliferation (Fig. 3C), potentially indicating the presence of the Wnt/STOP pathway in *Drosophila*^48^. Cluster IV reflected a set of genes that were strongly upregulated by Wg++ or armΔN and was enriched for chitin metabolism (Fig. 3D). As expected this cluster was enriched for binding sites of known targets of canonical Wnt signalling, including *ap* and *ind* ^49^ (Fig. 3E).

**Figure 3:**
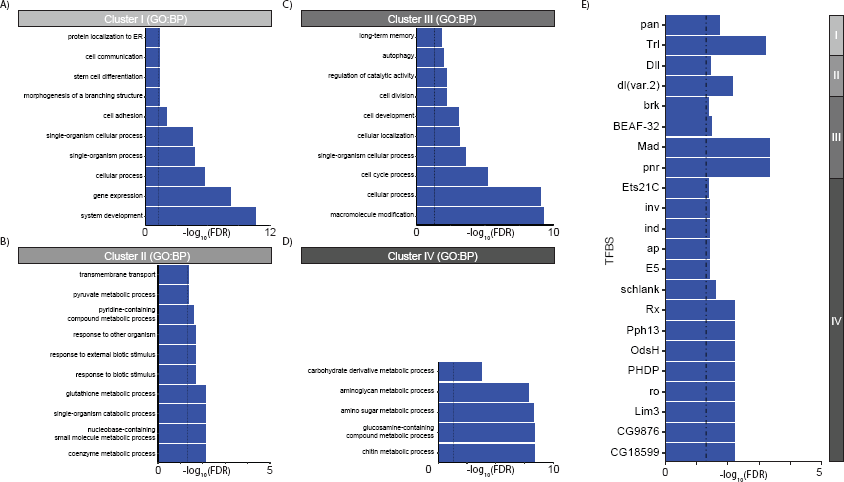
Functional annotation of clusters of genes with different responses to perturbing Wnt signalling. (A) Gene ontology (GO) enrichment for cluster I (B) GO enrichment for cluster II (C) GO enrichment for cluster III (D) GO enrichment for cluster IV. Where applicable ten representative GO terms (FDR < 5%) are displayed to summarise the enrichment profile for each cluster (see Methods). (E). TFBS enrichment (FDR < 10%) of each cluster.

Several studies have found that canonical and non-canonical signalling exhibit antagonistic effects on each other ^17,50,51^. Wg++ and Wnt4++ correspond to non-endogenous overexpression capable of driving these distinct antagonistic components of the Wnt pathway. Comparing expression of Wg++ against Wnt4++, we identified 1,798 genes as differentially expressed (Fig. 4A, absolute fold change > 1.5, FDR < 10%), with up- and down-regulated genes appearing to be associated with distinct biological processes and functions. Genes upregulated in Wg++ compared to Wnt4++ were associated with processes associated with cellular growth or the cell cycle, whilst those showing decreased expression were linked to cell adhesion, polarity and morphogenesis (Fig. 4B). Investigation of the promoter sequences of genes upregulated by Wnt4 compared to Wg revealed enrichment for binding sites of important developmental genes (i.e., *Ubx* and *cad*) (Fig. 4C). These results support the antagonism of canonical and non-canonical Wnt signalling and show Wg and Wnt4 as regulating vastly different downstream transcriptional programs.

**Figure 4:**
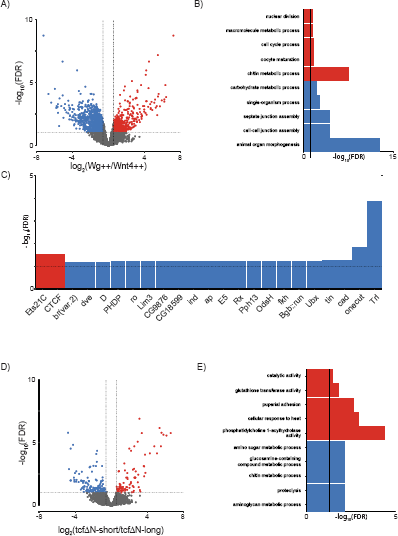
Functional annotation of clusters of genes with different responses to perturbing Wnt signalling. (A) Volcano plot of differentially expressed genes identified between Wg++ and Wnt4++. Significantly differentially expressed genes (absolute fold change > 1.5, FDR < 10%) are highlighted in red or blue for upregulated and downregulated genes respectively. (B) GO enrichments and, (C) TFBS enrichments for genes identified as upregulated (red) and downregulated (blue) between Wg++ and Wnt4++ implicates Wg and Wnt4 as driving different downstream transcriptional programs. (D) Volcano plot of differentially expressed genes identified between tcfΔN-short and tcfΔN-long. Significantly differentially expressed genes are (absolute fold change > 1.5, FDR < 10%) are highlighted in red or blue for upregulated and downregulated genes respectively. (D) GO enrichments for upregulated (red) and downregulated (blue) between tcfΔN-long and tcfΔN-short implicates an upregulation of genes involved in glutathione metabolism and stress in tcfΔN-short.

Despite the differences in the size of tcfΔN-short and tcfΔN-long embryos (Fig. 1), their expression profiles appeared to be highly similar (Fig. 2A). Surprisingly, only 222 genes were identified as differentially expressed between these conditions (Fig. 4D, absolute fold change > 1.5, FDR < 10%). Genes upregulated in tcfΔN-short were enriched glutathione transferase activity genes (i.e., *GstD4*, *GstD3*, *GstD9* and *GstD6*) and included genes known to be upregulated in response to severe stress (i.e., *TotA*, *TotC* and *TotX*)^52^, whereas the set of genes downregulated was enriched for proteolysis and chitin metabolism functions (Fig. 4E). These results suggest an upregulation of stress response and GSH depletion/redox state as a potential mechanism responsible for the differences in the sizes of tcfΔN-short and tcfΔN-long embryos.

### TCF occupancy confirmed at potential targets by HA-ChIP-qPCR

For those genes whose expression changed in response to perturbations in canonical Wnt-signalling, we investigated publicly available TCF ChIP-seq data from *Drosophila* embryos ^53^. As canonical Wnt-signalling primarily signals via TCF/pan, we expected to see an enrichment of genes bound by TCF in several sets of differentially expressed genes (i.e., Wg++, tcfΔN-short or tcfΔN-long). However, we observed that only those genes upregulated by Wg++ were enriched for TCF binding. Both technical (e.g., antibody specificity) and biological factors (e.g. differences between whole embryos and our naïve system) could potentially explain this lack of enrichment for TCF binding.

The simple embryonic system we have developed allows HA-tagged isoforms of factors of interest to be introduced into the system, making it possible to perform ChIP experiments against chromatin binding factors that either lack or only have low quality antibodies. We randomly selected 23 genes whose expression profile mirrored the size changes observed for tcfΔN-short, tcfΔN-long and Wg++ (Fig.5 A,B) (cluster IV). We performed ChIP-qPCR on their promoters from embryos expressing HA-tagged tcfΔN-long or HA-tagged tcfΔN-short constructs (Fig. 5C), to investigate whether these genes are direct or indirect target of Wnt signalling in this system. In addition, we investigated whether H3K27me3, a histone mark associated with repressed and bivalent genes ^54^, was present at this set of gene promoters (Fig. 5D). We identified two genes (CG13806 and CG7252) whose promoters were bound by tcfΔN-short-HA and showed high levels of H3K27me3 in tcfΔN-short embryos, suggesting that these genes are direct targets of Wnt/β-catenin repression in our system. Peritrophin-15A was found to lack tcfΔN-short-HA at its promoter but showed high H3K27me3 signal, suggesting that this gene is an indirect target of Wnt/β-catenin repression, whose expression is regulated, at least in part, via repressive histone modifications.

**Figure 5:**
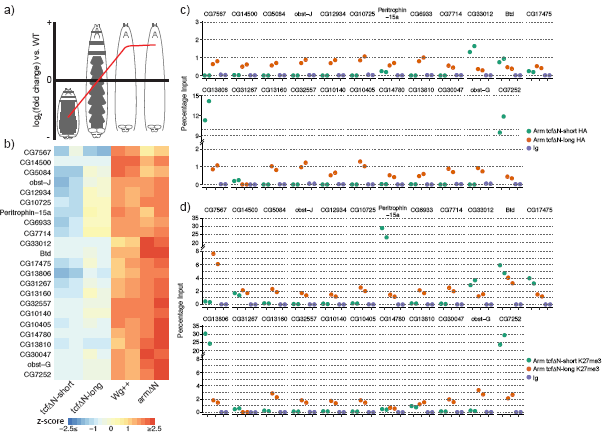
HA-ChIP followed by qPCR in embryonic system identifies putative targets of Wnt/TCF mediated repression. (A) Cluster IV is contains a series genes that showed strong upregulation in Wg++ and armΔN, mild downregulation in tcfΔN-long, and strong downregulation in tcfΔN-short indicating that these genes are under regulation by canonical Wnt signalling. (B) Expression profiles of 23 randomly selected genes from cluster IV (C) Promoter occupancy by HA ChIP of binding of tcfΔN-short and tcfΔN-long at selected promoters, and (D) H3K27me3 enrichment at selected promoters in tcfΔN-short and tcfΔN-long identifies CG13806 and CG7252 as genes which are potentially directly repressed by Wnt/TCF signaling.

### Blocking Apoptosis restores cells to epidermis

The differences in size between the tcfΔN-long and tcfΔN-short can be explained at least in part by two processes, either the embryo is growing less or cells are undergoing more apoptosis. Our transcriptional analysis identified an upregulation of stress response in tcfΔN-short embryos (Fig. 4C, D), which could implicate either of the processes. To evaluate how expression of tcfΔN-long or tcfΔN-short influences cell division and apoptosis in our system, we performed immunostaining using anti-phospho-H3 antibody and anti-cleaved caspase 3, respectively. Immunostaining was performed on the embryos collected from flies overexpressing Wg and used as control in the experiment. As apoptosis begins at stage 11-12 during *Drosophila* embryogenesis ^55^, we chose stage 14 embryos for immunostaining. We could not detect much apoptosis at these stages, but we did observe a large number of cell divisions occurring in wild type and Wg++ embryos (Fig. 6 C, D, G, H). There was a small increase in apoptosis in tcfΔN-short as quantified in Fig. 6I.

**Figure 6:**
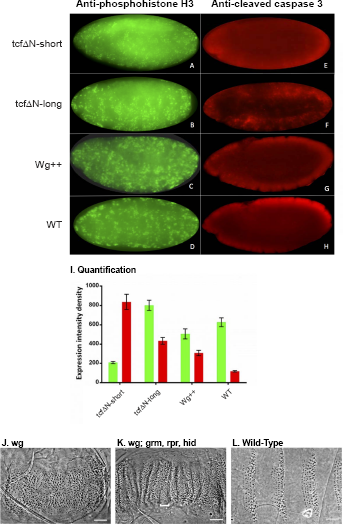
Comparison of cell division and apoptosis markers in developing embryos. (A-H) Embryos of the three canonical Wnt signaling conditions stained for the cell division marker phosphohistone H3 and apoptosis marker cleaved caspase3 as compared to wildtype. (I) Quantification of fluorescence from images of immune-stained embryos. (J-L) Blocking apoptosis rescues some cells in the small embryo phenotype of *wg* mutants by the H99 deletion.

These differences might indicate increased apoptosis as a potential mechanism to explain the smaller embryos in tcfΔN-short. Therefore, we tested whether blocking apoptosis could rescue the embryo size phenotype. To block apoptosis in small embryos, we used a deletion (Df(3L)H99) which removes three apoptosis genes *grim*, *reaper*, and *hid* ^56^. We combined H99 with a *wg* mutation and observed a restoration of cells between the segment boundaries (Fig. 6K, note the small denticles, bracket) as compared to *wg* mutant alone (Fig. 6J) and WT (Fig. 6L). This was similar to previous finding using *arm* mutants along with H99 ^57^. Although some of the embryo length is restored in *wg*; H99 double mutant embryos, the embryos do not resemble the large embryos of tcfΔN-long (Fig. 1), suggesting that apoptosis alone cannot completely explain the effect of tcfΔN-short.

### Discussion

We show that our naïve embryo system is amenable to quantitative analysis of both whole-body phenotype and associated transcriptional responses to perturbations targeting specific components and branches of the Wnt signalling pathway. The Wnt signalling pathway is of particular interest for this type of analysis as it consists of a canonical pathway with a well-defined mechanism of signal transduction, and a series of cell polarity pathways regulating a variety of cellular behaviours ^13,58^. These pathways can be thought of as a signalling network ^59,60^, where upon signal activation a poorly defined mechanism selects the pathway and outcome. Our system allowed the observation of specific transcriptional profiles that were clustered depending on which branch and which component was perturbed, illustrating clear differences in the sets of regulated genes and the involved biological processes.

For the canonical pathway, much but not all of the cellular response is mediated through β-catenin and TCF transcriptional activation and repression ^48^. For a strong activation of canonical Wnt signaling, we used overexpression of Wg, which resulted in full embryo growth (aside from a head involution defect ^55^). Phenotypically embryos generated by overexpression of Wg appear the same as those with a gain of function Arm allele. However, we found a marked difference between the two conditions with a large number of genes activated by Wg++ but not by armΔN (cluster III, Fig. 2A). These genes were associated with cell proliferation and the cell cycle. The results from our transcriptional analysis therefore suggest that Arm independent transcriptional activation occurs downstream of Wg, perhaps through the proposed Wnt/Stop mechanism ^47,48,53^, a pathway not previously documented in *Drosophila*.

For the strong loss of signaling condition, we performed two experiments using tcfΔN-short alone and tcfΔN-short along with armΔN. We observed that the transcriptional profile was very similar illustrating that all Arm dependent transcription requires a form of TCF that can interact with Arm (Fig. 2B). This re-establishes the Arm/TCF interaction as the main source of transcriptional activity due to Arm transactivation^5^. For the intermediate loss of signaling condition, we expressed a tcfΔN-long construct, a condition where we observe a loss of patterning, with most epidermal cells producing denticles but without a strong effect on embryo size. The identical phenotype was produced by the dominant negative DisArmed allele ^23^. As this is a highly-expressed form of the Arm protein that is immune to standard ubiquitin-mediated degradation and fails to act in transcriptional activation, the most likely explanation is that DisArmed binds to TCF, either sequestering it or preventing TCF from taking part in transcription. Either way, it perfectly phenocopies the absence of TCF (Fig. 1), and shows a very similar transcriptional profile to tcfΔN-long especially in clusters III and IV (Fig. 2A).

The fourth condition was the use of a non-canonical Wnt4 molecule that signals through a different receptor (PTK7/Otk) opposing Wg. We found that in all three conditions Otk expression, Wnt4 expression, and Wnt4/Otk expression a similar cohort of genes was regulated (Fig. 2A), and that the highly-correlated expression profiles of Otk and Wnt4 compared to WT support that they reside within the same section of the Wnt signalling pathway (Fig. 2C). The set of genes upregulated by perturbing Wnt4/Otk did not correspond to those upregulated by the canonical Wg pathway, and instead represent a new gene set involved in morphogenesis, cell:cell communication and adhesion (Fig. 3, 4), a finding that is in keeping with and correlates with the polarity pathways that determine cell shape and organization during epidermal development ^61-71^.

Recently, Wang and colleagues found that redox state in germ line stem cells was regulated by Wnt signaling ^72^. Different cellular states (i.e. proliferation, apoptosis, differentiation) have been associated with different redox states ^73,74^. Our transcriptional analysis indicated that glutathione metabolism, a metabolic pathway associated with regulation of redox potential in the cell, was regulated by Wnt signaling in our system. tcfΔN-short embryos showed an increase in the expression of genes associated with both glutathione metabolism and response to stress, whereas such a similarly strong change in expression was not observed in tcfΔN-long and Wg++. This finding suggests that embryonic Wnt signaling is required to modulate redox metabolism and its dysregulation in tcfΔN-short might result in increased stress and apoptosis ^75^. However, inhibition of apoptosis did not completely rescue the size differences between tcfΔN-long and tcfΔN-short suggesting the involvement of additional pathways.

Our gene expression and ChIP data do not support a simple explanation for which genes are activated and which are repressed. Previous studies attempting transcriptional profiling of genes downstream of Wnt have found a wide range of results and thousands of genes ^14,34,76^. For example, an early study looking at developmentally important transcription factors in *Drosophila* embryonic development found more than 1,000 sites where TCF was bound by ChIP-Chip ^34^. Since these genes are not expressed in the same way in different cells, it is likely that a complex combinatorial system with multiple transcription factors or epigenetic regulation is in place.

Overall, we present a useful *in vivo Drosophila* system that allowed us to characterize and bring together several aspects of Wnt signaling. We have looked at transcriptional repression and activation, moderate and strong canonical signaling conditions, and at the effects of opposing Wnt ligands. Showing the utility of our experimental system, our transcriptional analysis led us to identify a novel, *Drosophila* β-catenin independent set of genes activated by overexpression of Wg and completely different gene cohorts downstream of Wnt4 and Wg. We envision that detailed cellular and molecular studies in this naïve embryo system will allow to identify and test specific transcription factors and binding sites, and to delineate different signaling outcomes from different perturbations of Wnt and other signaling pathways.

## Supplemental Figures

**S1.**
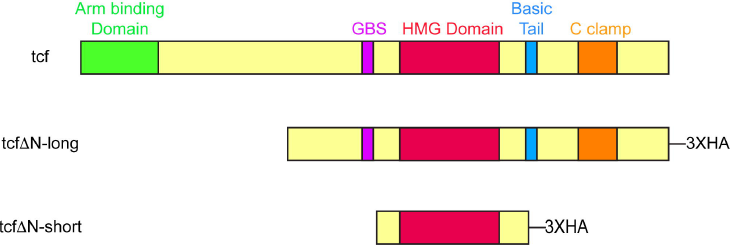
Domain structure of tcf and tcf alleles used in this study. The schema is adapted from ^15^ showing the Arm or β-catenin binding domain (green), the Groucho binding sequence (GBS, purple), the high mobility group domain (HMG, red), basic tail (blue), and C clamp (orange) regions. The deleted domains as well as the HA tags are represented for the tcf constructs generated for this study.

**S2.**
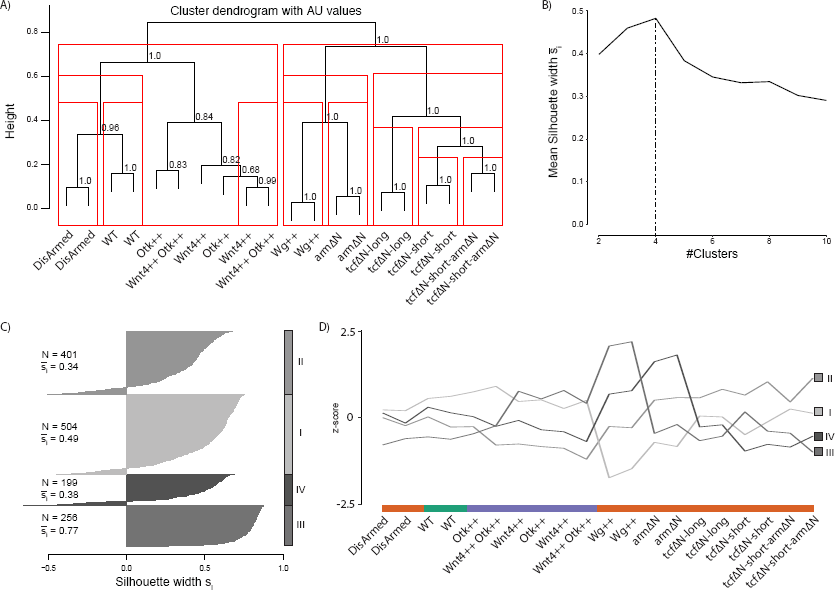
Assessment of clustering expression profiles in response to perturbing the Wnt signalling pathway. (A) Stability of observed sample clustering (Fig. 2A) was assessed by randomly bootstrapping the data (see Methods). Support for each branch over 10000 iterations is indicated and with a red box is present to highlight that the observed clustering is not expected by chance (AU>0.95). (B) The average silhouette width was calculated across different numbers of clusters to determine the optimal choice for the number of clusters (N=4). (C) Distribution of silhouette widths across all clusters. (D) Average expression profile (z-score) for each cluster over all conditions.

## Supplemental Data

Table S1: Excel sheet with results of the differential expression analysis.

Table S2: Excel sheet containing significant enrichments for sets of differentially expressed genes

Table S3: Excel sheet with Chip qPCR primer sequences

Movie S1: Movie of *bicoid*, *nanos*, *torsolike* embryos in phase contrast

Movie S2: Movie of lightsheet microscope using Ubi-NLS-GFP and UAS-myr-Tomato^77^ as nuclear and membrane markers respectively.

## Materials and Methods

*Fly strains and transgenics:* The *bicoid, nanos, torsolike* strain (*bcd*^*E*1^, *nos*^*L*7^, *tsl*^146^) ^39^ was recombined with DaGal4 flies to make a triple mutant with Gal4 driver. UAS-Otk-3XHA^19^, UAS-Wnt4^78^, UAS-DisArmed^23^, UAS-arm-ΔN^41,79-81^ were described previously.

1. *wg*^*IG22*^ ; Df (3L) H99
2. *bcd*^*E*1^, *nos*^*L*7^, *tslcd*^146^, da-Gal4 females x UAS-tcfΔN-long-3XHA
3. *bcd*^*E*1^, *nos*^*L*7^, *tslcd*^146^, da-Gal4 females x UAS-tcfΔN-short-3XHA
4. *bcd*^*E*1^, *nos*^*L*7^, *tslcd*^146^, da-Gal4 females x UAS-Wg
5. *bcd*^*E*1^, *nos*^*L*7^, *tslcd*^146^, da-Gal4 females x UAS-DisArmed
6. *bcd*^*E*1^, *nos*^*L*7^, *tslcd*^146^, da-Gal4 females x UAS-Wnt4
7. *bcd*^*E*1^, *nos*^*L*7^, *tslcd*^146^, da-Gal4 females x UAS-Otk
8. *bcd*^*E*1^, *nos*^*L*7^, *tslcd*^146^, da-Gal4 females x UAS-Wnt4, UAS-Otk
9. *bcd*^*E*1^, *nos*^*L*7^, *tslcd*^146^, da-Gal4 females x UAS-tcfΔN-short-3XHA, UAS-ΔN-Arm
10. *bcd*^*E*1^, *nos*^*L*7^, *tslcd*^146^, da-Gal4 females x UAS-arm-ΔN
11. *bcd*^*E*1^, *nos*^*L*7^, *tslcd*^146^, da-Gal4 females x Ubi-NLS-GFP; UAS-myr-Tomato

The TCF transgenes were made by PCR amplification of DNA from an ovarian library using primers:

TCF Short FOR—CACCATGGTTTCTGGAATTTTCGGGCTAAGTCAA

TCF Short REV—CGTTGTCGATCTGTCTTTTTTTCGCTTTTT

TCF Long FOR—CACCATGGCATTAGCTGCTATAGCACTGTCTAAT

TCF Long REV—TGAAACGCTAATAACGCCGTTATCGGAAGA

The PCR products were cloned into pENTR vectors (Invitrogen) and recombined using Gateway technology (Invitrogen) into pUASg.AttB.3XHA vectors for fly injection ^82^. The DNA was injected into strain P[CaryP]attP2 68A4 by BestGene Inc California ^83^.

### Microarray

We collected approximately 50 embryos per microarray experiment, with the control embryos being non-expressing naïve embryos. Extracted mRNA from the various genetic conditions was then analysed on Affymetrix Drosophila 2 microarrays by standard procedures. For each condition, we performed two biological replicates. Microarrays were normalised using GC-RMA. Prior to differential expression analysis, probesets were filtered by 1) removing probesets not mapping to a gene 2) removing probesets which mapped to multiple genes 3) if a gene had multiple probesets assigned to it the probeset with the largest IQR was used 4) probesets not showing expression greater than 2.5 in at least two samples and those mapping to non-canonical chromosomes were removed. Following these preprocessing steps, differential expression analysis was performed using LIMMA ^48^. Clustering of gene expression profiles was performed by converting gene expression to z-scores and clustering them using (1 - cor) / 2 as a dissimilarity measure. To assess the stability of sample level clustering we used pvclust with 10000 iterations to calculate approximately unbiased (AU) p-values (Figure S2A). All of the major expected clusterings remained stable. The set of samples relating to perturbation of the non-canonical pathway (Wnt4++, Otk++, Wnt4Otk++) did not form a stable cluster but were highly unstable between each other. The optimum number of clusters to cut the gene-associated dendrogram was determined by calculating the mean silhouette width over a number of different cluster sizes (Fig. S2B,C).

Enrichment for Gene ontology was performed using a hypergeometric test from the GOStats package^84^. Results from GO enrichments were simplified for presentation purposes by filtering terms with a high semantic similarity^85^, all significant results (adjusted p-value < 0.05) from the enrichment analyses are available in Supplemental Table S2. Enrichment for pathways was performed using ReactomePA^86^, all significant results (adjusted p-value < 0.05) are available in Supplemental Table S2. For pairwise comparisons, enrichments performed using the set of genes used in the differential expression analysis as the background, whereas for the enrichments based on the clustering (Fig. 2A) the background was the set of genes identified as differentially expressed over all conditions.

Enrichment for TFBS motifs was performed using AME (from the MEME suite^87^) against the JASPAR 2016 database^88^. Promoter regions were defined as 1kb upstream/downstream of a gene’s Ensembl-annotated TSS (dm6, Ensembl version 86). p-values were corrected using fdr, with a TFBS classified as significant at an FDR of 10%.

### ChIP-seq analysis

ChIP-seq data for *TCF* at embryonic 0h-8h and 16h-24h was downloaded from modENCODE and lifted over from dm5 to dm6 ^53^. A gene was defined as been bound by *TCF* if there was at least one identifiable peak within 2kb of the gene’s Ensembl-annotated TSS (dm6, Ensembl version 86). A hypergeometric test was used to calculate if TCF binding was overrepresented in defined sets of genes.

### Embryo Collection and immunostainings

Embryos were collected 14-16 hrs after egg deposition and dechorionated with bleach and fixed with 4% formaldehyde in presence of heptane and sodium phosphate buffer and vortexed at maximum speed. Embryos were devitellinized in methanol/heptane and stored at -20c until needed. Immunostainings were performed by standard methods with respective antibodies and Alexa Fluor dyes ^67-69^. Whole embryo images were taken under 20X magnification with Zeiss Axiocam. Quantification of fluorescence was done using ImageJ software tools ^89^. FFT band pass filter was applied to the images for correction of any uneven illumination and horizontal scan lines acquired by phase contrast microscope followed by conversion to 40 pixels. For fluorescence quantification in the cells, small structure default pixels were optimized to 3 pixels and tolerance threshold was set at 5% using binary process function. Intensity density was obtained by using the particle analyser tool. Standard error was calculated using data from n=3 for each condition and error bars were plotted.

### Antibodies

The following antibodies were used in the study: polyclonal Anti-phosphorylated histone H3 (Millipore, #06-570) and Cleaved caspase 3 (Cell signaling Technology, #9661) for embryo staining as cell division marker and apoptosis marker respectively. Hoechst stain was used to image nuclei (Invitrogen). Rabbit polyclonal to HA tag (Abcam, #ab9110) antibody, mouse monoclonal (mAbcam6002) to Histone H3 (tri methyl K27) and Anti RNA polymerase II (Millipore #05-623B) were used in chromatin immunoprecipitation experiments.

### Chromatin Immunoprecipitation (ChIP)

Embryos staged around 14-16hrs were collected and cross linked with 1.8% formaldehyde in presence of heptane. Cell and nuclear lysis was done by respective lysis buffers (Easy Magna Chip kit (Millipore)) and Wheaton Dounce homogenizer was used to achieve uniform lysis. Chromatin was sheared for 18 cycles (30sec ON and 30Sec OFF) by sonication (Diagenode Bioruptor®) to a size range of 200bp -1kb chromatin fragments and the size was checked on a 2% agarose gel. Anti-HA tag (Abcam, #ab9110) and H3K27me3 (Abcam, #6002) antibodies have been used to immunoprecipitate the DNA. Chromatin was diluted 10 fold in Chip dilution buffer, control sample was saved and immune complexes were prepared and incubated at 4ºC overnight with respective antibody and protein A/G magnetic beads (Millipore). Subsequent washing of immune complexes was performed with low salt, high salt, LiCl immune complex wash and TE buffer, and then eluted in Elution Buffer. After reverse cross-linking and Proteinase K treatment, ChIP and control DNA samples were prepared and purified with columns (Millipore). IgG and IgM were used as negative controls in the ChIP assay.

### Real-Time PCR

RT qPCR primers set which can amplify 150–200 base pair fragments were designed (NCBI primer design tool) for the 23 short listed genes for evaluating ChIP assays from the indicated genomic regions. Realtime PCR was carried in a PikoReal96 Real Time PCR system (Thermo Scientific) following the manufacturer’s instructions. Gene-specific transcription levels were determined in a 10 μl reaction volume in triplicate using QuantiFast SYBR Green and qPCR was conducted at 95°C for 7 min, followed by 40 cycles of 95°C for 5s and 60°C for 1 min. Two biological replicates were used to perform the experiment and results have been replicated. The specificity of the reaction was verified by melt curve analysis. Primer sequences are available in Supplemental Data 3.

## Data availability

All microarray data from this study is available from GEO under accession number GSE97873.

## Acknowledgements

We thank Xiaoping Liu who made the original TCF constructs. We thank Eric Wieschaus and Ken Cadigan for fly strains, and J. Bischof and K. Basler for constructs. This work was supported by an Academic Research Fund (AcRF) grant (MOE2014-T2-2-039) from the Ministry of Education, Singapore to N. Tolwinski.

## Contributions

J. S., K. K. L., P. K., J. J., J. B. L., N. H., N. S. T. performed the experiments. N. H., E. P., and N. T. wrote the paper. All authors reviewed manuscript.

## Competing interests

The authors declare no competing financial interests.

